# A flexible framework for automated STED super-resolution microscopy

**DOI:** 10.1101/2025.05.05.652196

**Authors:** David Hörl

## Abstract

Super-resolution microscopy enables the observation of cells at unprecedented detail but usually entails high light exposure and slow imaging. Thus, often only a few manually selected regions are imaged, limiting the ability to capture the distribution of quantitative features in a population of cells in an unbiased fashion. An exciting strategy to circumvent these limitations are imaging pipelines in which informative regions are detected on-the-fly by software and imaged automatically. Point-scanning methods like STimulated Emission Depletion (STED), in particular, can be sped up by selective imaging of small regions. Here, I present autoSTED, a flexible Python-based framework to construct automated imaging pipelines for STED microscopy. Instead of fixed acquisition loops defined at the beginning of an experiment, autoSTED employs a priority queue of acquisition tasks. After each image acquisition, callback functions can trigger actions like adding new tasks based on data, enabling dynamic and adaptive imaging. Complex experimental pipelines can be built from easily exchangeable building-blocks or expanded through custom code, facilitating integration of state-of-the-art computer vision methods. autoSTED can drastically speed up super-resolved imaging of subcellular structures and enables autonomous operation of a microscope for days with minimal hands-on time and bias.

## Introduction

Light microscopy has been the prototypic single-cell technique for centuries and remains invaluable for studying the spatial organization and dynamics of cells. Many cellular phenotypes, from the expression levels of proteins^1^ to their subcellular spatial localization^2^, show a large variability from cell to cell. Thus, a large number of cells must be observed to accurately capture the complex distribution of quantitative features within a population. Historically, microscopy has often been an anecdotal method with results relying on a few hand-selected images. In recent decades, developments in electronic control of instruments and digital image acquisition have opened the way to higher throughput in microscopy^3^. Still, microscopy remains limited due to long measurement times and can produce large amounts of data that often require specialized analysis tools and dedicated storage strategies to handle^4^. Super-resolution microscopy^5^ suffers from these limitations in particular, as it trades enhanced resolution for slower acquisitions. Techniques such as single molecule localization microscopy (SMLM) or structured illumination microscopy (SIM) rely on multiple exposures, not only lengthening measurement times but also producing large datasets that need to be processed into the final images. Postprocessing-free techniques such as STimulated Emission Depletion (STED) don’t produce large amounts of data but rely on slow point scanning. Thus, super-resolution microscopy often remains limited in the number of biological entities studied in an experiment. Hardware methods of improving the throughput of super-resolution microscopy remain an active area of research, e.g. through parallel scanning in STED or RESOLFT^6–9^. As faster acquisitions usually have lower signal-to-noise ratio (SNR), software approaches like image restoration^10^, i.e. using postprocessing to reconstruct high SNR data from low SNR input, have gained popularity. Crucially, in many biological samples only a fraction of the specimen may be of relevance to the research question at hand. Modern, computer-controlled microscopes raise the possibility of combining hardware and software approaches by automating the microscope for selective imaging. In such approaches, image acquisition is coupled to real-time data analysis to only acquire data at informative locations.

The potential of automated microscopy has been apparent for over a decade and several impressive proofs-of-principle have been implemented over the years, like the seminal early example of the Micropilot framework^11^ of Conradt and colleagues, that implemented hierarchical overview-detail imaging pipelines on a variety of microscopes. In addition to manufacturer-supplied software interfaces, the MicroManager project^12,13^ has emerged as an open-source solution for microscope control and formed the basis for more advanced automated imaging strategies^14^. The rise in popularity of the Python programming language led to new microscope control frameworks being implemented in this language^15–19^ as well as the establishment of interfaces between Python and MicroManager^20^. As a promising strategy to overcome several drawbacks of super-resolution microscopy, automated imaging has been combined with STED^21–23^, SMLM^24–26^ and SIM^27^. Automated imaging can be especially beneficial to point-scanning techniques like STED that, while generally slow, allow for dynamic switching of the scanned field-of-view (FOV) and pixel size. In addition to selective acquisition of relevant data, smart microscopy can also be reactive and adjust parameters during a live-cell acquisition based on the sample^28^ or even interact with cellular state through optogenetics or microfluidics^16,29,30^. Finally, like in many other areas, deep learning has emerged as a powerful tool to use in microscope automation, facilitating the detection of regions or events of interest^27,29,31^ to image or on-the-fly image restoration^32^ beyond the capabilities of classical methods.

Here, I present autoSTED, a modular and open-source computational framework for the automation of STED microscopy written in Python. Instead of pre-defined acquisition loops tailored to specific experiments, autoSTED employs a priority queue of individual acquisition tasks. After each image has been acquired, a set of callback functions are run which can add new tasks to the queue but may also be used to perform other actions such as sending notifications to users via email or waiting for interactive feedback. Due to the flexible and modular design, autoSTED allows the construction of complex imaging pipelines from a library of reusable building blocks and thus adaptation to a variety of tasks such as hierarchical imaging, systematic search for rare phenotypes or adaptive change of parameters during time series imaging. It abstracts hardware specifics and thus makes it easy to include custom code, e.g. for deep learning-based detection of objects of interest. On the following pages, I will highlight the design principles behind my framework as well as novel imaging strategies that can be implemented using this foundation.

## Results

### A framework for flexible image acquisition pipeline design

The autoSTED framework works with commercially available Abberior Instruments microscopes and builds on the foundation offered by the microscope control software Imspector and its Python interface through the SpecPy library, which allows accessing and setting microscope parameters (laser powers, stage or scan positions, …) through parameter trees represented as Python dictionaries. This interface, combined with the vast ecosystem of Python libraries for image processing^33^, machine learning^34^ and general scientific computing^35^ makes it easy to develop simple, linear pipelines to automate specific imaging tasks. I went one step further and aimed to implement autoSTED as a generalized and flexible framework.

At the heart of an autoSTED acquisition pipeline lies a dynamic priority queue of parameters dictionaries with which images should be acquired (here called *acquisition tasks)*. The main loop of an acquisition pipeline run will pick the highest-priority task from this queue, run the acquisition and save resulting data. Here, the priority is not only given by the order in which tasks were added, but also their hierarchy level, e.g., a super-resolution detail image will have high priority and be imaged before moving to the next confocal overview image. For each hierarchy level, a set of callbacks can be registered that are invoked after an image has been acquired. These callbacks typically take the form of *acquisition task generators* that may add new acquisition tasks to the queue. After all callbacks have been processed, the main loop continues with the next acquisition task from the queue (Figure 1A).

**Figure 1:**
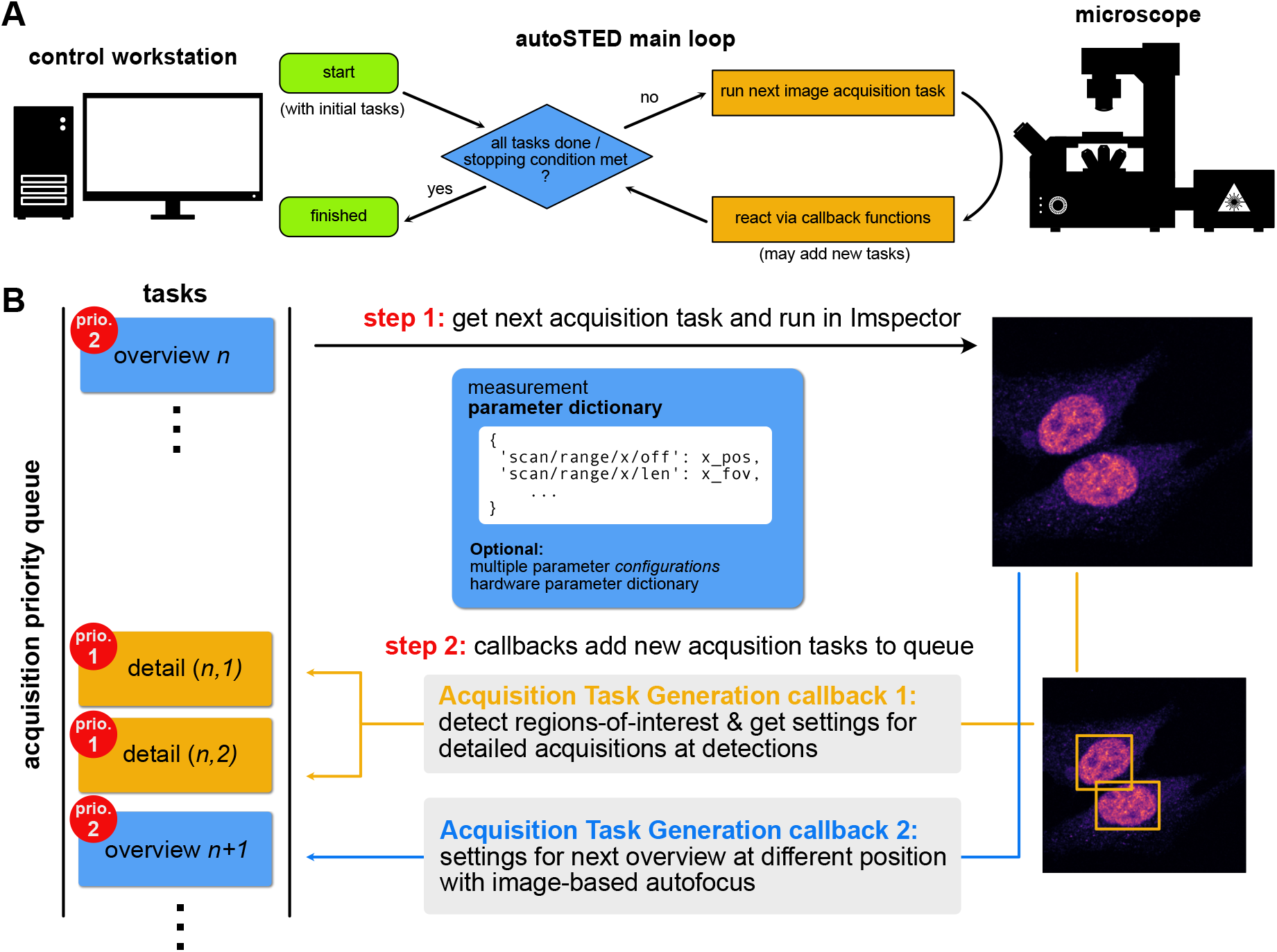
A: Flowchart of the main control loop of autoSTED. B: Schematic overview of one iteration in an automated imaging run using a modular autoSTED pipeline. The main acquisition loop will get the next acquisition task (wrapping measurement parameter dictionaries) from a priority queue and acquire an image based on those parameters. After the measurement, a set of callback functions attached to the hierarchy level of the measurement are called to, e.g., enqueue acquisition tasks for detailed measurements of ROIs within the just imaged field-of-view (FOV) and enqueue the next overview measurement.

An example of an iteration of the main loop for overview-detail hierarchical imaging is shown in (Figure 1B). Here, the pipeline has a task for an overview image (priority 2) in the queue (top). After an overview image has been acquired with these parameters, the callbacks associated with overview images are invoked: The first callback analyzes the acquired overview image, searches for cells and for each detected cell places acquisition tasks for detail images at those locations with (higher) priority 1 on the queue. A second callback adds the next overview task to the queue. This way, overview tasks are iteratively added to the queue instead of all being defined at the beginning of the experiment. This makes it possible to keep the pipeline running for an indefinite amount of time and to perform on-the-fly image-based adjustments of overview parameters (e.g. focus position) and thus facilitate robust imaging for hours. After handling all callbacks, the high-priority detail tasks will be imaged in the main loop before moving on to the next (lower priority) overview image. The pipeline will keep running until either there are no more tasks in the queue, or a user-defined stopping criterion is met (maximum imaging time, desired number of images).

The acquisition task generation callbacks should produce complete parameter sets that define an image to be acquired. However, in practice only a fraction of parameters, mainly stage positions or scan offsets, change from image to image, while parameters like pixel sizes or laser powers (usually) remain constant for each image of a given hierarchy level. For flexibility and reusability, I added the possibility to assemble top-level acquisition task generation callbacks (producing full parameter sets) from smaller building block callbacks (producing a subset of parameters) that are grouped in an AcquisitionTaskGenerator object. For example, the callback for enqueueing detail images (callback 1 in Figure 1B) may consist of distinct steps (illustrated in Figure 2A): 1) loading basic settings that don’t change, e.g. laser powers, from a file 2) taking the stage position of the last overview image and 3) detecting objects in the last overview image and generating scan offsets and ranges for them, specifying the regions-of-interest (ROIs) to be imaged. For the object detection in step 3), autoSTED provides generic wrappers for user-defined functions that accept images as NumPy arrays and return pixel coordinates, bounding boxes of objects or segmentation maps. The wrappers translate pixel-level results to the microscope’s coordinate system and parameters. That way, own or third-party object detection code (e.g. cell detection via Cellpose^36^, Supplementary Note 2.4) can easily be integrated and adjusted for a given experimental task while leaving interactions with the microscope hardware as-is. Some of the building blocks produce only a single set of parameters (e.g. the general settings of step 1), whereas others may produce multiple (e.g. scan positions for each object detected in step 3). Thus, after the individual building-block callbacks are run, the AcquisitionTaskGenerator will automatically broadcast these parameters (repeating single parameter sets similar to NumPy^35^) and enqueue an acquisition task with complete parameters for each combination (Figure 2B).

**Figure 2:**
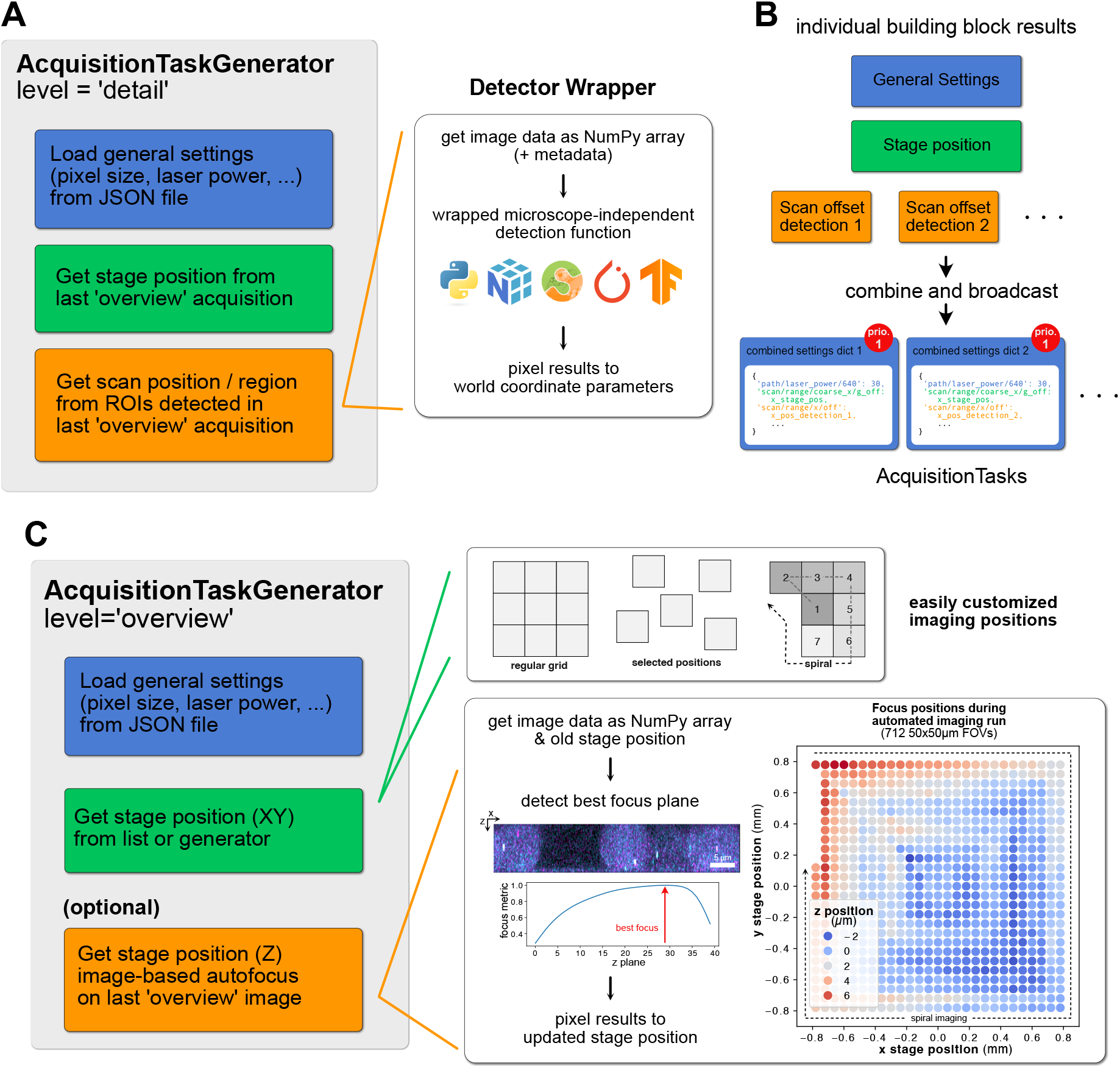
Building acquisition task generation callbacks from simple, reusable building blocks. A: individual blocks of a callback to enqueue detail images as shown in (Figure 1B). It will generate acquisition tasks by combining general settings from a file, stage position parameters of the previous overview image and scan offsets defined by a spot detector on the previous overview image. The spot detector can be further split into steps like loading data, performing detection using microscope-independent code (with the possibility to use a wide array of Python libraries) and translating pixel results back to scan positions for the generated acquisition task. B: Results of the individual building blocks are combined into complete parameter dictionaries by the AcquisitionTaskGenerator. The resulting tasks will be added to the acquisition task queue at priority level “detail”. C: Building Blocks of the callback to enqueue the next overview images as shown in (Figure 1B), with customizable locations and updates to the stage focus position based on the previous overview. The image-based autofocusing step can again be decomposed into individual steps. Right: Example of updated stage positions at 712 fields-of-view during an autonomous acquisition from^37^, showing effects like thermal drift (concentric fluctuations around center) or uneven slides (top-left to bottom-right gradient) that are compensated.

As a second example, the callback for enqueuing the next overview stack (callback 2 in Figure 1B) also consists of multiple steps that are grouped in an AcquisitionTaskGenerator object: It will 1) load basic settings from a file, like for the detail images, 2) set the stage position based on the next position in a predefined list or generator and 3) change the z-drive position to keep the sample in focus based on the previous overview stack (Figure 2C). Here, autoSTED again offers simple customizability by e.g. changing the stage positions to image at. They can be in a regular grid, or a list manually selected by the user. Likewise in this flexible framework, optional steps, like the autofocus step can be added if needed or removed if a hardware focus-lock is being used instead, for example.

By iteratively generating new positions to image at, e.g. in a growing spiral, the pipeline can keep the system running indefinitely, with the queue of acquisition tasks never emptying. Thus, after each image acquisition in the pipeline, stopping criterions, also implemented as callbacks, are checked to interrupt the pipeline. Pre-built stopping criteria are provided to halt the pipeline after a target number of images has been acquired or a specified time has passed. Again, stopping criteria objects are just thin wrappers around Python functions returning a Boolean “stop/don’t stop” decision, allowing for easy extension.

A detailed user guide of autoSTED is available as Supplementary Note 2.

### autoSTED facilitates DNA-FISH data acquisition at kilobase-resolution

I primarily developed the autoSTED framework for studies of the spatial arrangement of small genomic loci using DNA fluorescense in-situ hybridization (FISH) (Figure 3A-C) in our group. The first application was in a study^37^ in which we performed DNA-FISH of arrays of ~5kb spaced genomic loci in both prototypically active and inactive chromatin. Likewise, we used the framework to showcase systematic single-locus DNA-FISH using optimized probe generation protocols^38^ and study promoter-enhancer dynamics during pluripotency exit^39^.

**Figure 3:**
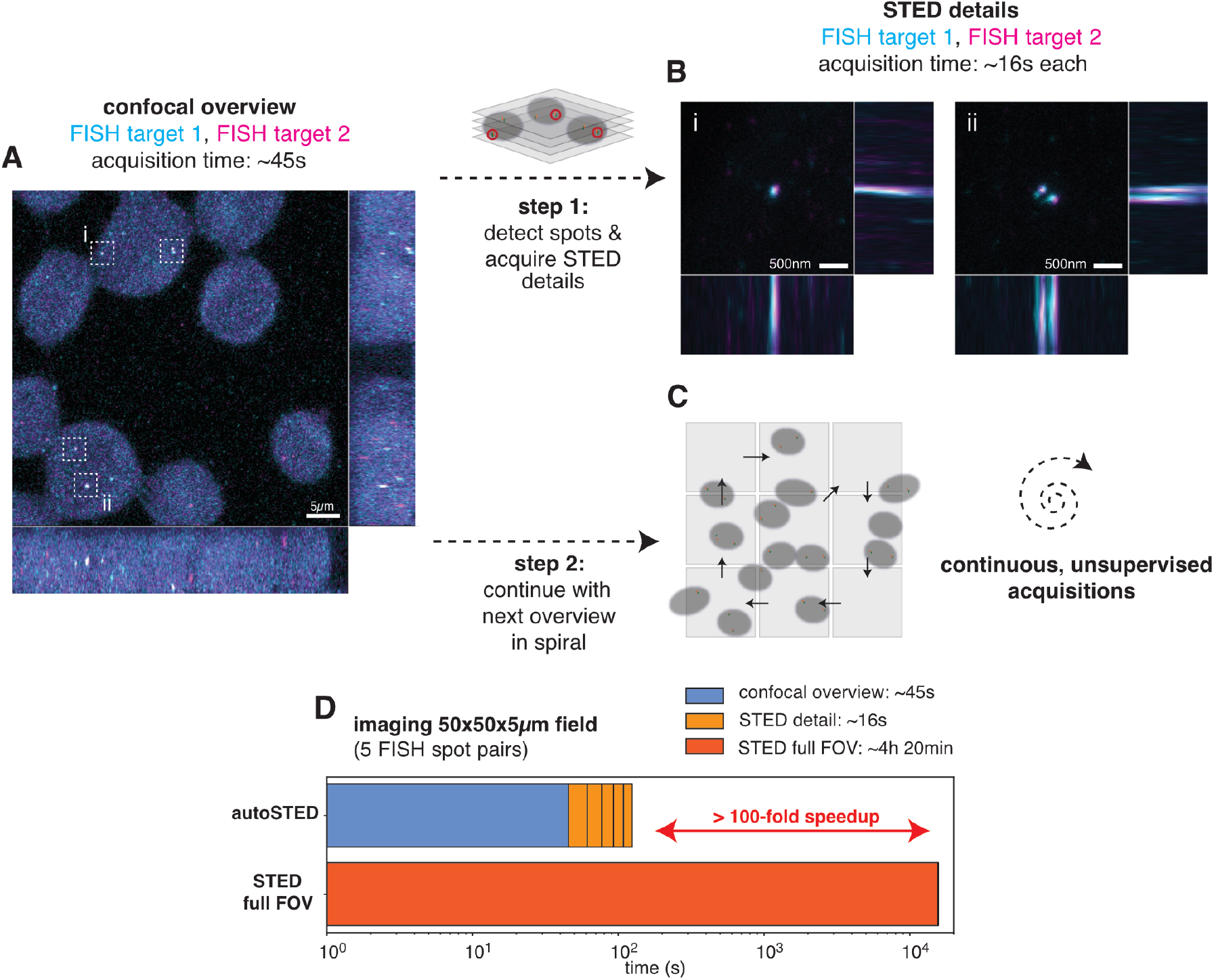
Schematic of the (confocal) overview-(STED) detail acquisition strategy used for imaging pairs of DNA-FISH spots. After the acquisition of a confocal overview stack (A: orthogonal maximum projections), pairs of signals (dashed boxes) are detected and small, detailed STED stacks (2D depletion pattern) are acquired around them (examples in B), allowing the resolution of closely spaced FISH spots in individual channels, e.g. due to freshly replicated loci (FOV ii). Acquisition times are based on parameters used in^37^. Afterwards, the pipeline continues with the next overview in spiral pattern, allowing for continuous imaging (C). D: Estimated time to image a whole FOV at STED resolution, compared to using autoSTED (confocal overview and 5 STED details) shows an over 100-fold speedup using my framework.

STED microscopy has several advantages for our applications. In addition to the high spatial resolution necessary to resolve multiple genomic elements labelled in one color (Figure 3B), chromatic errors between channels are drastically reduced when measuring in several color channels^40^. Furthermore, in contrast to stochastic super resolution techniques, STED microscopy offers the possibility to switch from fast confocal mode to super-resolved STED mode only at informative loci. This advantage becomes clear in a simple approximative calculation based on the imaging experiments in^37^: Imaging a whole 50×50×5µm FOV at STED pixel sizes and dwell times would take over 4 hours and lead to premature bleaching of faint FISH signals. In contrast, imaging the whole field at lower resolution and dwell times takes only 45 seconds, followed by super-resolution imaging of small fields (3×3×1.4µm) at 16 seconds each, a roughly 100-fold speedup when assuming 5 ROIs per overview FOV (Figure 3D). To make use of this potential speedup, an automated pipeline that detects ROIs to image on its own, as offered by autoSTED, is necessary. Using an overview-detail pipeline as described above, we could thus let our system acquire data autonomously overnight, resulting in several hundreds of super-resolved images per sample.

### On-the-fly stitching facilitates automatic imaging of large objects

Small subcellular structures like organelles or FISH spots can be readily imaged using a simple overview-detail strategy as outlined above. However, when trying to detect larger objects, such as whole cells or nuclei, the exact ROI to image in super-resolution often can’t be determined from one image alone as cells lie on the border of overview images (Figure 4A). Individual images can’t be made arbitrarily large as the FOV of the high numerical aperture (NA) objectives used in STED is limited. Common solutions are acquiring overview images with a low magnification/NA but high-FOV objective and then switch to a high NA objective for detailed acquisitions. Keeping focus, handling of immersion oil and the danger of collisions makes fully automated imaging using this strategy challenging. Another solution is to image a large overview comprised of tiled images first, stitch the tiles and detect objects to image in the stitched overview. Since acquiring many images may take considerable time, sample drift can become a problem here as cells from earlier images may no longer lie at their observed position. This can be mitigated by registering images relative to the latest acquisition, but the issue remains as drift may continue during the subsequent detailed imaging of many detected cells.

**Figure 4:**
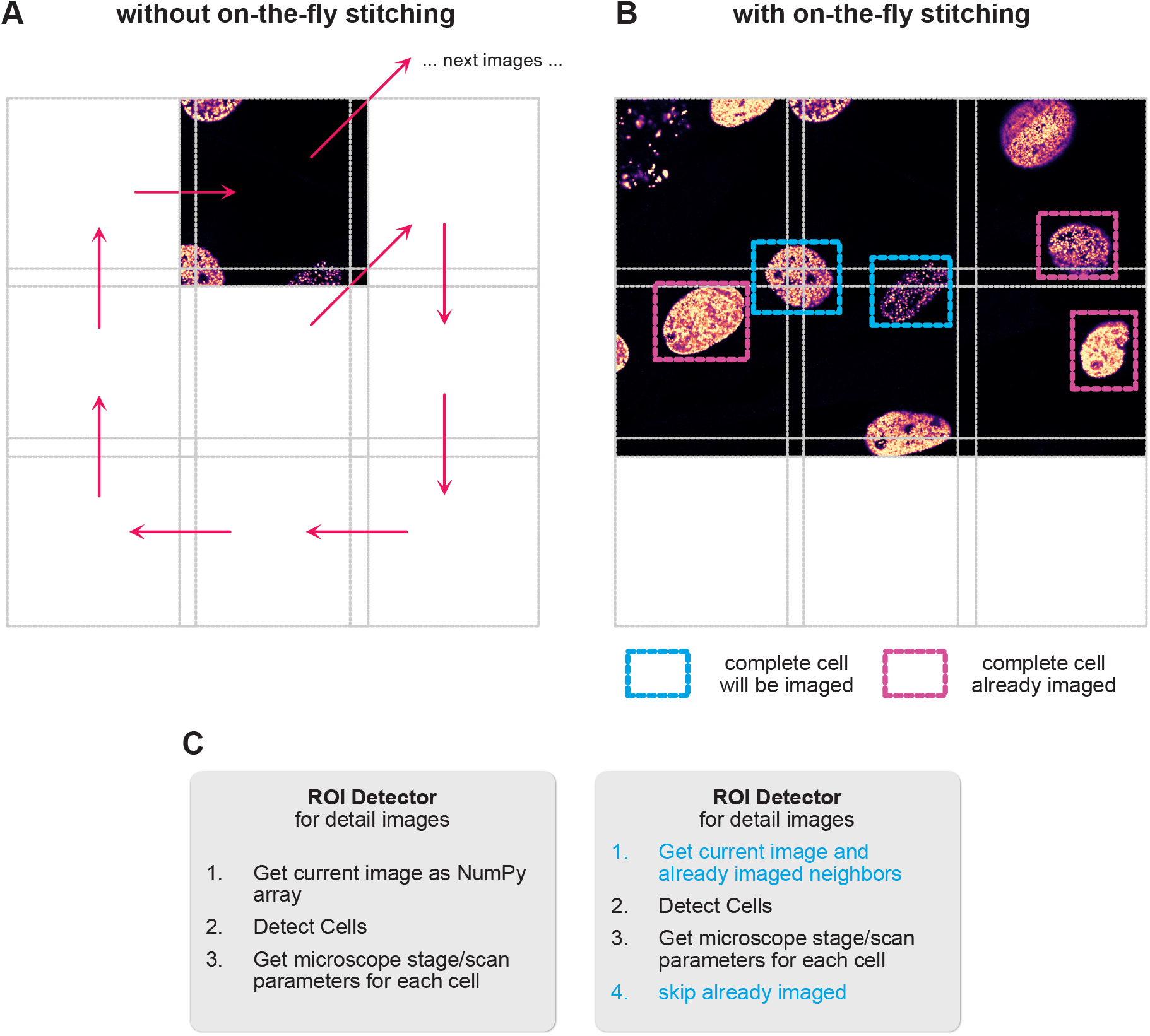
Schematic of on-the-fly stitching of overview images. A: When detecting whole cells to image in detail in individual overview images, cells may lie on the border of the image and have to be skipped. B: With on-the-fly stitching, the detection callback will not only consider the last overview image, but also stitch it with previously acquired neighboring tiles, allowing the detection of cells in border regions that are not completely present in any one individual image. C: Due to the flexible, building block-based framework, this change only requires replacement/addition of some steps of the callback, while others can be left as-is.

Instead, I decided to implement an on-the-fly stitching strategy. In short, when a callback accesses the latest overview image to detect objects in it, a specialized data selector building block also collects overlapping overviews imaged earlier and stitches them to the current one. The stitching can be done based solely on stage/scan coordinates but also with an extra registration step placing all tiles relative to the current one, thus mitigating drift effects.

Again, I took advantage of the modular nature of the framework, in which the detector callback is built from several steps: 1) getting the most recent image data as a NumPy array, 2) detecting pixel-level bounding boxes around objects of interest and 3) translating the pixel coordinates back to a parameter dictionary of stage/scanner positions. To enable on-the-fly stitching, I replaced just the first step responsible for loading the most recent overview with one that stitches all surrounding tiles on-the-fly, with all other parts being re-used (Figure 4C). To prevent multiple imaging of the same object, I also added a check to discard already imaged ROIs.

This strategy saw application when we applied autoSTED in a study on the induction of senescence-like phenotypes in cells using the small molecule inflachromene (ICM)^41^. Here, we supplemented biochemical and omics data with super-resolution imaging data of DNA-stained nuclei. Senescent cells often have enlarged nuclei, exacerbating the problem of cells lying on the border between overview tiles. With on-the-fly stitching, we could easily acquire hundreds of super-resolution images of nuclei of both young, senescent, and ICM-treated cells to quantitatively study their global chromatin texture.

### Easy construction of complex imaging pipelines

The applications highlighted above employed two-step overview-detail imaging pipelines in which regions of interest were detected in confocal overview stacks and selectively imaged at higher resolution (Figure 5A, top). This already provided a drastic speedup over naïve imaging at the highest resolution, but the flexible nature of autoSTED makes it easy to build multi-step pipelines to further increase imaging efficiency. For example, one can introduce an additional pre-scan step before each overview in which a very quick and low-resolution image is acquired. A corresponding callback would only enqueue a proper overview if at least some signal is detected in the pre-scan (Figure 5A, middle). That way, sparse samples can be quickly traversed and FOVs containing no cells can be skipped with minimal time overhead. Another option is to introduce an intermediate step between the overview and (subcellular) detail levels. One could, for example, detect cells in the overview image and then take images of each cell before detecting and imaging subcellular details in those cells (Figure 5A, bottom). This way, overview stacks can have reduced z extent (or overviews can be just a single plane), whereas larger z-stacks are acquired at the cell level, similar to^23^. My framework also allows construction of more complex experiments, e.g., multiple detection callbacks can be used to detect different types of cells or objects in overview images and acquire detail images of them with different parameters (e.g. different laser combinations depending on whether cells express a marker or not).

**Figure 5:**
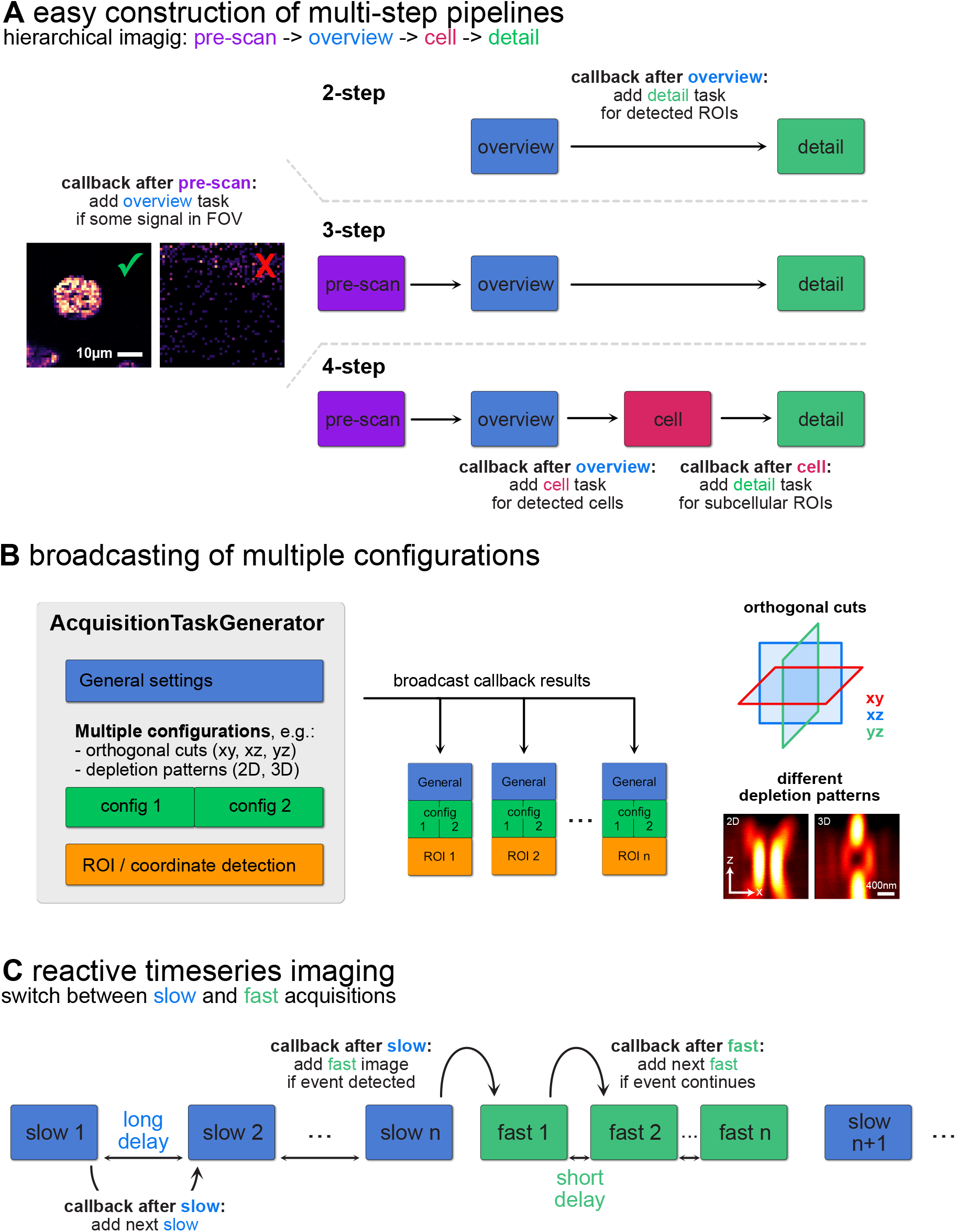
Options for creating complex imaging pipelines in autoSTED. A: Schematic flowcharts of multi-step hierarchical image acquisition pipelines. Aside from two-level overview-detail strategies (top), the framework allows adding different levels, such as a fast pre-scan (middle) or intermediate cell-level (bottom). B: By providing multiple configurations in a building block of an AcquisitionTaskGenerator, the settings will be broadcast to enable, e.g., imaging each detected ROI with both 2D and 3D depletion patterns. C: Tasks generated by callbacks can have a delay which can be used to implement reactive timeseries imaging by switching between slow and fast acquisition rates.

Another way of building more complex pipelines comes from the possibility of creating acquisition tasks with multiple configurations. Using the broadcasting logic of the AcquisitionTaskGenerator objects described above, one can acquire multiple images for each detected object/coordinate of interest. This way, one can, e.g., image the same structure with different STED depletion patterns or acquire just orthogonal slices at a location instead of a full stack (Figure 5B).

Finally, although the system was built with hierarchical imaging of fixed samples in mind, reactive timelapse imaging similar to^21,27^ can also be implemented via a delay that can be added to acquisition tasks. Using a two-level hierarchy, one can acquire repeating slow “search” images with long delay until a callback enqueues higher-priority, low delay “event” images that are repeated a set number of times or until the event of interest is no longer detected (Figure 5C).

## Discussion

Modern quantitative biology requires the observation of numerous cells to accurately capture the variability inherent in biological phenomena or discover rare phenotypes^42^. For example, promoter-enhancer interactions during transcription initiation^43^ may only last a short time but still have lasting effects on the state of a cell. Microscopy readily provides cellular and subcellular resolution but scaling up to high throughput is challenging, especially when focusing on fine details that require high or even super-resolution microscopy. To overcome this limitation, many have turned to automating microscopes^11^. Attempts to make microscope automation available in a user-friendly way include the hardware-agnostic graphical interface MicroManager^12–14^ or interfaces to the popular Python programming language^15,17,20^. In this context I present autoSTED as a Python-based framework for automated STED nanoscopy. It reconciles user-friendliness and flexibility by allowing the easy construction of automation pipelines from existing building blocks while also offering adaptability to many tasks through Lego-like combination of the blocks and various customization hooks. The framework was successfully employed in multiple studies in the field of chromatin biology in our group over several years^37–39,41^ in which it was used to automate repetitive tasks and allowed the collection of data for hours to days without user input.

My automation strategy is based on a dynamic priority queue of acquisition tasks and building block-based callbacks to add tasks to this queue. I chose this architecture for its flexibility: multiple hierarchy levels in an acquisition can thus be handled without constructing nested loops and pipelines can be adapted to new experimental needs by exchanging individual callbacks or building blocks without a hard-to-maintain rewrite or copy of the whole pipeline.

Callback-based frameworks have been used for automated microscopy to various degrees^20,26,29,44^. Conceptually, the SMLM-focused PYME platform of Barentine and colleagues^26^ shares many similarities with my work. Their framework also represents an experiment as a priority queue of acquisition tasks, with the ability to put new ROIs to image on the queue as a result of analysis callbacks called recipes, that can be built from smaller building blocks. The large amounts of data involved in an SMLM experiment require a complex software architecture though, e.g. distributing work across a compute cluster. In contrast to this, the deterministic postprocessing-free and hardware-based super-resolution offered by STED is a unique advantage in the context of automated imaging, as the amounts of data recorded are relatively small, allowing direct processing of raw data on the microscope control workstation. Furthermore, as acquisition time in point-scanning methods like STED scales with the size of FOVs imaged, selective acquisition of small regions provides an immediate speed benefit for these methods.

Hiding complicated hardware details behind a software abstraction is a common task in software development, ranging from operating system-wide device drivers^45^ to scientific instrument control as done by MicroManager. In this spirit, my framework provides pre-built wrappers to generate acquisition tasks from pure Python functions that perform coordinate or object detection on raw pixels in the form of NumPy arrays. The wrappers automatically translate pixel-level results to microscope parameters like physical stage or scan coordinates.

Thus, users familiar with common Python libraries for image processing can adapt the framework to their needs without having to worry about the minutiae of the hardware. That way, state-of-the-art computer vision approaches like deep learning-based segmentation methods^36,46^ can be easily integrated to detect objects to image in an automated microscopy pipeline. The flexible nature of autoSTED also allows for easy incorporation of techniques such as software autofocus or on-the-fly stitching to facilitate long-term autonomous imaging via drop-in callback building blocks. Furthermore, due to the extensibility with pure Python functions, users can also employ common inter-process communication strategies included directly in the Python standard library or third-party modules to, e.g., perform object detection on a remote compute server using Web APIs, control other hardware like microfluidics systems or send status reports to the experimenter via email.

The framework builds upon the SpecPy interface that has been successfully used to implement automated imaging pipelines focused on specific tasks^19,22,23,32^. An impressive example showcasing the strengths of smart STED microscopy is the event-triggered STED (etSTED) pipeline of Alvelid and colleagues^21^. In this work, the authors employed a custom-built microscope allowing for both camera-based widefield imaging and STED. They constantly monitored samples in widefield mode and detected events of interest (spikes in calcium or other fluorescent indicators, vesicle fusion) via on-the-fly image analysis. Upon detection of an event the system switched imaging modes, and a short time series in STED mode was imaged at the region of interest. The small ROI allowed this system to acquire high framerate super-resolution data of synapse and vesicle dynamics. Complementary to bespoke solutions employing custom-built hardware, autoSTED seeks to offer a flexible framework to enable various automation tasks on microscopy setups controlled through Imspector.

autoSTED helps in overcoming two main drawbacks of STED microscopy: slow speed and photobleaching/-toxicity resulting from high laser intensities. It has been shown that both can be ameliorated with selective imaging of small regions^47^. However, manual selection of regions to image can be a time-consuming process, especially for faint punctuate signals like in DNA-FISH. It is, however, a comparatively easy task in image processing and can be automated using my framework. Adaptive illumination strategies^48,49^ can also prevent photobleaching in such a situation but do not provide a timing advantage per se. If it is supported by the microscope, adaptive illumination can be easily integrated into the framework, as relevant parameters can be set via the SpecPy interface. Furthermore, due to the reactive nature of autoSTED, these parameters can also be set adaptively. This way, parameters like DyMIN thresholds for a detail acquisition can be based on information from overview scans instead of being statically set to a fixed value for all images.

The implementation presented here is specific to hardware from one manufacturer, but the general strategy of queue- and callback-based modular microscope automation is a powerful approach applicable to many complex imaging experiments. Due to its modular design, autoSTED could also be adapted to other systems with relative ease by replacing a few interface classes. As a proof-of-principle, autoSTED allows for simulated microscopy in pre-recorded datasets instead of an actual microscope for demonstration purposes, which is implemented by replacing just one interface object. For integrating other microscopes, the main functionality that needs to be provided is getting and setting microscope parameters based on parameter dictionaries (acquisition tasks), acquiring data with the current state and accessing data as NumPy arrays. One example of microscope control software that would allow these interactions with minimal effort is MicroManager through the Pycro-Manager interface^20^.

For quantitative biology, automated microscopy as offered by autoSTED is a crucial step in the direction of more reproducible data. It can minimize hands-on time of scientists and replace the bias of hand-picking with an objective decision of which cells to image and thus enable autonomous recording of more representative data for days.

## Materials and Methods

### Available fundamentals

The image acquisition workflow offered by the manufacturer’s software, Imspector, is simple: most of the global microscope parameters (calibrations, etc.) as well as parameters of an individual measurement (stage position, scan area, laser powers, etc.) can be set via text fields in the graphical user interface. Once the parameters are set by the user an acquisition can be started, resulting in the acquisition of one or more image stacks. The main units of an experiment in the software are measurements, which can consist of multiple configurations. Each configuration can have its own measurement parameters as well as acquired image data stacks. All data of one measurement are by default saved as one file in the MSR format. To facilitate work with MSR files in Python, I implemented a reader library (Supplementary Note 1).

The software is complemented with the SpecPy^50^ Python interface that exposes the majority of this streamlined workflow via simple function calls: new measurements and configurations can be created, their parameters can be accessed and modified in in the form of Python dictionaries. Acquisition of the currently active measurement can be started, and the resulting image data stacks can be accessed and modified in the form of NumPy-arrays.

### Microscope hardware

I developed and tested autoSTED using an Abberior Instruments (Expert line) STED microscope based on an Olympus IX83 body with 100x (NA 1.4, oil immersion), 60x (NA 1.2, water immersion), 20x (NA 0.75) and 10x (NA 0.4) objectives (all UPlanSApo, Olympus), pulsed 488nm, 594nm and 640nm excitation lasers and a 775nm pulsed depletion laser, a galvanometric scanner and APD detectors for each excitation line. Phase modulation to generate depletion patterns is done via a SLM-based easy3D module (Abberior).

### Microscope control workstation

The microscope is controlled via a PC integrated in the main electronics rack, equipped with an AMD Ryzen 7800X3D CPU, a NVIDIA GeForce RTX 4070 GPU and 64 GB RAM running Windows 10.

### Software environment

The development of autoSTED happened over several years with changing versions of Imspector (the current major version 16.3 and earlier). I currently work with Imspector 16.3.19720 (2024.08), SpecPy 1.2.3 and Python 3.11. I also tested the code on a new demo system (Infinity) provided by the manufacturer and encountered no problems. The framework should therefore be usable on any relatively recent Abberior STED setup controlled through Imspector.

## Data availability

The FISH data showcased here (Figure 3) are taken from a dataset published with a previous study and are available under https://osf.io/zjwxm. The focus map in Figure 2C is based on acquisitions of slide A209 from this dataset.

Example overview images in Figures 4, 5, S1 show newly replicated DNA in C2C12 mouse fibroblasts labelled with Abberior STAR635P conjugated via a click-reaction to EdU incorporated during a 10 min pulse as described in^51^. A demo dataset of the same sample is available under https://osf.io/et4br/?view_only=23d060ed9a8f48b6863852ed65ac2c45.

## Code availability

The code of the autoSTED automation framework and example pipelines are available under a MIT license at https://github.com/hoerlteam/sted_automation. The current version at the time of publication of this preprint is tagged as version v2.0.0 and is also available under https://doi.org/10.5281/zenodo.15287446.

Code for the msr-reader library is available under a MIT license under: https://github.com/hoerlteam/msr-reader. It is also available via PyPI (https://pypi.org/project/msr-reader/) and can thus be installed easily via pip.

## Acknowledgements

I thank Pascal Bawidamann for help in implementing the first prototype version of the autoSTED framework. I thank the users of the framework (Katharina Brandstetter, Simona Nasiscionyte, Clemens Steinek, Gabriela Stumberger and Josefin Sedlmeier) for providing data and constructive feedback. I thank Hartmann Harz and Tobias Ragoczy for the initial planning of the project and Heinrich Leonhardt for constant support and feedback.

This work was supported by a research grant from Deutsche Forschungsgemeinschaft (DFG, grant number HO 7333/1-1).

## Notes

https://osf.io/et4br/?view_only=23d060ed9a8f48b6863852ed65ac2c45

https://osf.io/zjwxm

### Competing Interest Statement

The authors have declared no competing interest.

