## Supplemental Notes for "A flexible framework for automated STED super-resolution microscopy"

#### Supplementary Note 1: A pure Python library for reading MSR files

Imspector saves acquired data in a format called MSR, which itself is a superset of a documented format called OBF [1]. The files can be read in Fiji [2] via the Bioformats [3] library. Several Python bindings to Bioformats exist, but they rely on complex interactions with Bioformats and Java under the hood and often make the setup of a computational environment challenging.

Thus, to streamline Python-based processing of acquired data, I implemented a pure Python library, called msr-reader [4], for reading pixel data and metadata from MSR/OBF files based on the format description and made it publicly available under the MIT license. The module is available to end users via pip and has no external dependencies other than NumPy, allowing it to be used with minimal requirements to the computational environment. In short, the library provides an OBFFile (reader) class, which users can instantiate given the path of a file. Typical use cases are illustrated in (Figure S1). After parsing file contents, a reader object provides easy access to image pixel data as NumPy arrays and common metadata such as pixel sizes, image shapes or stack names. Furthermore, it also provides access to the full microscope metadata saved to files as XML strings either following the SpecPy parameter format or standardized OME-XML. These metadata can be parsed further with a third-party XML library, e.g. the ElementTree module provided by the Python standard library or the ome-types [5] library.

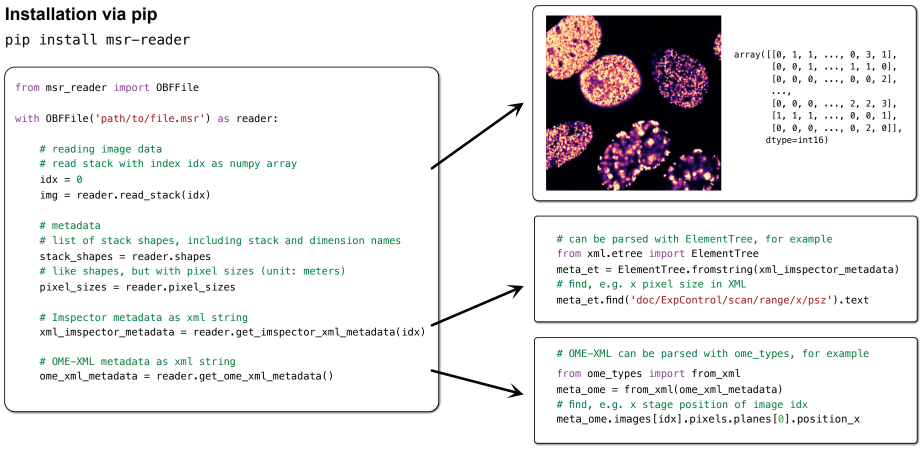


Figure S1: Example usage of the msr-reader library. Upon opening a file with an OBFFile reader, users can access pixel data of individual stacks as NumPy arrays, common metadata like pixel sizes or even the complete Imspector or OME metadata of an acquisition in XML format, which can be further parsed using third-party XML libraries.

### Supplementary Note 2: autoSTED User Guide

This short user guide is a compilation of install instructions and usage example notebooks (basics, parameter saving and overview-detail imaging of cells) from https://github.com/hoerlteam/sted_automation. Have a look there to get interactive Jupyter notebook versions of this guide, for further examples and up-to-date instructions.

##### Warning

**autoSTED is designed to automate operation of expensive scientific equipment. While we don't do anything that is not also possible manually via Imspector or SpecPy, be careful to avoid damage to the microscope (e.g. through collisions of stage and objective) or danger to yourself (e.g. from lasers).**

- **SpecPy and therefore autoSTED uses SI units, so a movement or image size of `1.0` would correspond to 1 meter. Stages and scanners should typically stop when they reach their limits, but nonetheless, pay attention to specify, e.g. distances in the appropriate fractions of meters (in Python you can use scientific notation like `5e-6` to indicate $5 * 10^{-6}$ (m) = 5 micron).**
- **Never use autoSTED to operate a microscope when laser safety measures are disabled!**

#### 2.1: Installation

##### SpecPy and compatible NumPy

Our framework talks to the microscope control software Imspector via the SpecPy interface.

SpecPy `.whl` files can be found in the Imspector installation folder (e.g. `C:\Imspector\Versions\{Ver_Nr}\python\specpy\{SpecPy_Ver_Nr}`). SpecPy is built for specific Python and NumPy versions, indicated in the file/folder name, so it is a good idea to create a conda environment with the corresponding versions (e.g. for Python 3.11.4, NumPy 1.24.3):

```
conda create -n autosted-env python=3.11.4 numpy=1.24.3
```

Then, in you new environment, you can install SpecPy:

```
conda activate autosted-env
pip install C:/Imspector/path/to/specpy.whl
```

##### autoSTED

Once you have an environment set up, you can install autoSTED:

```
git clone https://github.com/hoerlteam/sted_automation.git
cd sted_automation
pip install .
```

Alternatively, install directly without cloning:

```
pip install git+https://github.com/hoerlteam/sted_automation.git
```

If you do not have git on your system, you can also download the code from the GitHub page https://github.com/hoerlteam/sted_automation manually, unzip and install via `pip install .` in the downloaded folder.

##### Aviod NumPy updates

**Warning:** Installing additional third-party packages may sometimes update NumPy, causing SpecPy to stop working. Consider using the `--dry-run` options of `pip/conda` to check if an installation might cause problems.

Sometimes, it helps to explicity re-state the NumPy version during install:

```
pip install package-of-choice numpy==1.24.3
conda install package-of-choice numpy=1.24.3
```

Pinning the NumPy version in `conda` might also be worth a look: https://docs.conda.io/projects/conda/en/latest/user-guide/tasks/manage-pkgs.html#preventing-packages-from-updating-pinning

#### 2.2: Basic autoSTED usage concepts

This section showcases how our **autoSTED** framework and its core components of an acquisition task queue with callbacks works in principle. It follows the `basics.ipynb` example.

We will first show what acquisition tasks are, how they can be added to the acquisition task queue by callbacks and how a pipeline can be built from small, reusable, building blocks. For more realistic applications, please also check out the other notebooks in the examples folder.

**Note:** Imspector should be open in the background.

**We also provide the `examples/demo_overview_detail.ipynb` example notebook that performs virtual microscopy in a pre-recorded dataset, allowing testing of autoSTED without a microscope. The concepts from this user guide also apply in those simulated acquisitions.**

The main class to run an acquisition is `AcquisitionPipeline`:

```
from autosted import AcquisitionPipeline

# activate logging, as autoSTED uses it for some output
import logging

logging.basicConfig(level=logging.INFO)
```

To create an `AcquisitionPipeline` instance, we have to give it a path to which data should be saved and a list of the *hierarchy levels* of the images to acquire.

Here, we make a pipeline with only one hierarchy level `'image'` for demonstration purposes, but in realistic scenarios this could be something like `['overview', 'detail']`, indicating that we first take overview images in which we then take detail images.

The acquisition can then be started with the `.run()` method:

```
pipeline = AcquisitionPipeline(
    data_save_path="acquisition_data/test",
    hierarchy_levels=["image"],
)

pipeline.run()
```

This will do nothing (except creating the output directory if it does not yet exist). That is because we have not added any *acquisition tasks* to the pipeline's queue.

##### Acquisition Tasks

Acquisition tasks are made from **parameter dictionaries** that correspond to the value trees used in Imspector.

For example, let's get the measurement parameters of the current measurement as well as the hardware parameters from Imspector via SpecPy:

```
import specpy as sp

# connect to Imspector
imspector = sp.get_application()
```

```python
# get current measurement parameters as dict
measurement_parameters = imspector.value_at(
    "", sp.ValueTree.Measurement
).get()
# get hardware parameters / calibrations as dict
hardware_parameters = imspector.value_at(
    "", sp.ValueTree.Hardware
).get()

measurement_parameters
```

An *acquisition task* usable in our pipeline is a list of pairs of measurement and hardware parameters:

```python
# make acquisition task:
# list of (measurement parameters, hardware parameters) pairs
acquisition_task = [
    (measurement_parameters, hardware_parameters)
]

# NOTE: you can use empty hardware parameters ({})
# (if you do not want to change them)
# unless you know what you are doing this may be the safer option
acquisition_task = [(measurement_parameters, {})]
```

We can add the task (at hierarchy level "image") to the queue of our pipeline using `.enqueue_task()`.

If we run the pipeline afterwards, the measurements in the queue will be run one after the other:

```python
pipeline = AcquisitionPipeline(
    data_save_path="acquisition_data/test",
    hierarchy_levels=["image"],
)

# enqueue our task
pipeline.enqueue_task("image", acquisition_task)

pipeline.run()
```

This will make a new measurement in Imspector, run an acquistion with the parameters we have specified, save it and then stop because we only added one task to the queue.

The resulting image data will be saved in the output path with filename: `{random string}_image_0.msr`.

Instead of the random prefix you can manually specify a prefix for your files via the `file_prefix` parameter of AcquisitionPipeline.

We can, of course, add multiple tasks to the queue:

```python
pipeline = AcquisitionPipeline(
    data_save_path="acquisition_data/test",
    hierarchy_levels=["image"],
)

# enqueue task twice
pipeline.enqueue_task("image", acquisition_task)
pipeline.enqueue_task("image", acquisition_task)

pipeline.run()
```

This will run the same measurement twice, resulting in two output files.

###### Advanced: Multiple Configurations

The reason why our Acquisition Tasks are a list of parameter pairs (instead of a single one) is that this way, we can support multiple configurations.

The following code will run the same acquisition twice, but as two configurations of one measurement:

```python
# measurement with two configurations
double_acquisition_task = [
    (measurement_parameters, {}),
    (measurement_parameters, {}),
]

pipeline = AcquisitionPipeline(
    data_save_path="acquisition_data/test",
    hierarchy_levels=["image"],
)

pipeline.enqueue_task("image", double_acquisition_task)

pipeline.run()
```

Here, the resulting data will be saved to one file, similar to manually making multiple configurations in Imspector.

###### Task Generation Callbacks

Instead of manually putting a list of measurements to be done in our pipeline's queue before running them all, we typically generate tasks via **callbacks**. This way, new tasks can be added in response to acquired data while the pipeline is running.

In principle, a task generation callback can be any function (or callable object) that returns a hierarchy level and a list of acquisition tasks (in the format specified above):

```python
def task_generation_callback():
    return "image", [acquisition_task]
```

```
task_generation_callback()
```

A callback can be run once at the beginning of the acquisition by passing it to the `.run()` method:

```
pipeline = AcquisitionPipeline(
    data_save_path="acquisition_data/test",
    hierarchy_levels=["image"],
)
pipeline.run(initial_callback=task_generation_callback)
```

More realistically, we might want to call the callback repeatedly after each image.

This can be done via the `.add_callback()` method of the pipeline, which will cause the callback to be run every time a measurement of a given level is finished.

**Note:** Since this keeps re-adding the acquisition task to the queue, it will run indefinitely until you manually stop via the stop button in Jupyter or by pressing `Ctrl-C`. In this case, the currently running image will be finished and the pipeline will stop afterwards.

```
pipeline = AcquisitionPipeline(
    data_save_path="acquisition_data/test",
    hierarchy_levels=["image"],
)

# add task_generation_callback
# to be run after each acquisition of level 'image'
pipeline.add_callback(
    task_generation_callback, level="image"
)

pipeline.run(initial_callback=task_generation_callback)
```

To not run forever, we can add *stopping conditions* to our pipeline. For example, a `MaximumAcquisitionsStoppingCriterion` will cause the pipeline to stop after a certain number of images have been acquired.

In the `autosted.stoppingcriteria` module, we also have stopping criteria to e.g. stop after a specific time.

```
from autosted.stoppingcriteria import (
    MaximumAcquisitionsStoppingCriterion,
)

pipeline = AcquisitionPipeline(
    data_save_path="acquisition_data/test",
    hierarchy_levels=["image"],
)

pipeline.add_callback(task_generation_callback, "image")
```

```
# add stopping criterion to stop after 5 images
pipeline.add_stopping_condition(
    MaximumAcquisitionsStoppingCriterion(5)
)

pipeline.run(initial_callback=task_generation_callback)
```

##### Assembling callbacks from building blocks

Until now, we have only run the same acquisition over and over. To actually do something useful, we want our callbacks to enqueue acquisitions with different parameters each time they are run.

For this, we offer a variety of *building block callbacks* in `autosted` that return a subset of parameters. E.g. a `SpiralOffsetGenerator` will generate new stage positions each time it is called:

```
from autosted.callback_buildingblocks import (
    SpiralOffsetGenerator,
)
from autosted.imspector import (
    get_current_stage_coords,
)

# generator of stage positions in a spiral
# with 50x50 micron steps, starting at current position
stage_position_generator = SpiralOffsetGenerator(
    move_size=[50e-6, 50e-6],
    start_position=get_current_stage_coords(),
)

# call 4 times and print result
for _ in range(4):
    print(stage_position_generator())
```

Multiple callbacks can be combined using an `AcquisitionTaskGenerator` object.

In the followng block, we use this to construct a combined callback that will first load full measurement parameters from a dictionary (or file if we instead give it a file path) using a `JSONSettingsLoader` and then overwrite just the stage position parameters with changing values supplied by the `SpiralOffsetGenerator`. Finally, the merged parameters will be returned as acquisition tasks at level "image."

```
from autosted.taskgeneration import (
    AcquisitionTaskGenerator,
)
from autosted.callback_buildingblocks import (
    JSONSettingsLoader,
)
```

```python
next_overview_generator = AcquisitionTaskGenerator(
    "image",
    # building block 1: return base measurement parameters
    JSONSettingsLoader(measurement_parameters),
    # building block 2: return stage coordinates
    # (moving in spiral every time it is called)
    SpiralOffsetGenerator(
        move_size=[50e-6, 50e-6],
        start_position=get_current_stage_coords(),
    ),
)

pipeline = AcquisitionPipeline(
    data_save_path="acquisition_data/test",
    hierarchy_levels=["image"],
)

pipeline.add_callback(next_overview_generator, "image")

pipeline.add_stopping_condition(
    MaximumAcquisitionsStoppingCriterion(5)
)

pipeline.run(initial_callback=next_overview_generator)
```

#### 2.3: Saving Imspector measurement parameters as JSON

We typically save parameters of a measurement to JSON text files, so we can re-use them in automated measurements using a `JSONSettingsLoader` callback.

While parameters can be set from the automation code we provide dedicated callbacks for some (e.g., pixel size, FOV length), in general, the idea is to save *template parameters* for all types of images you wish to acquire in an automation run (overviews, STED details, . . . ) and only change things like stage/scan position dynamically during the run. The saved parameters can be loaded with a `JSONSettingsLoader` as described above.

##### Hints

- it's best to deactivate easyCommander before saving parameters
- pay attention to settings like "lock aspect ratio" in Imspector (making image size settable in only one dimension)
    - if a setting is grayed-out in the GUI, we can't change it from code either

##### Basic saving

We want to save the whole Imspector parameter tree, which we can get as a dictionary via the `.value_at('', ...)` method.

Then, we can just save it using the `json` module:

```
import json

import specpy as sp


# get current measurement parameters and hardware parameters
imspector = sp.get_application()
current_measurement_parameters = imspector.value_at(
    "", sp.ValueTree.Measurement
).get()
hardware_parameters = imspector.value_at(
    "", sp.ValueTree.Hardware
).get()

# dump to JSON
with open("parameters.json", "w") as fd:
    json.dump(current_measurement_parameters, fd, indent=1)
```

#### 2.4: Overview-Detail imaging with autoSTED

Let's start with what we had at the end of section 2.2, running overview acquisitions in a spiral.

Here, we added a few small changes:

- instead of using the parameters of the current measurement, we use parameters saved to a JSON file as shown above.
- we wrap the settings loader in a `LocationRemover`: this just removes any location-related parameters (i.e. stage/scan offsets)
  - this is not strictly necessary here, as those parameters will be overwritten by the `SpiralOffsetGenerator`, but to keep everything clean, it still makes sense.

**Note:** When building your own pipeline, you could also build upon the other `overview_*` notebooks in the examples folder for:

- imaging in a regular grid (optionally with on-the-fly stitching)
- imaging at manually picked locations
- image-based autofocus in the overviews
- selective overview imaging with a pre-scan

```
from autosted import AcquisitionPipeline
from autosted.taskgeneration import AcquisitionTaskGenerator
from autosted.callback_buildingblocks import (
    JSONSettingsLoader,
    LocationRemover,
    SpiralOffsetGenerator,
)
from autosted.imspector import get_current_stage_coords
from autosted.stoppingcriteria import (
```

```python
    MaximumAcquisitionsStoppingCriterion,
)

import logging

logging.basicConfig(level=logging.INFO)


# pipeline with a single overview level: "image"
pipeline = AcquisitionPipeline(
    data_save_path="acquisition_data/test",
    hierarchy_levels=["image"],
)

# path to parameters saved as JSON
overview_config = "config_json/test_overview.json"

# overview generator combines settings from file
# with next stage positions in spiral
next_overview_generator = AcquisitionTaskGenerator(
    "image",
    LocationRemover(JSONSettingsLoader(overview_config)),
    SpiralOffsetGenerator(
        move_size=[50e-6, 50e-6],
        start_position=get_current_stage_coords(),
    ),
)

# add the callback and a stopping condition
pipeline.add_callback(next_overview_generator, "image")
pipeline.add_stopping_condition(
    MaximumAcquisitionsStoppingCriterion(5)
)

pipeline.run(initial_callback=next_overview_generator)
```

##### Accessing the data of a run

An `AcquisitionPipeline` stores acquired data (and the parameters used) in its `.data` attribute, which acts like a `dict`.

The keys are tuples of (str, ints) pairs, consisting of the level name and running count for each hierarchy level. E.g. the first overview image has index `(("image", 0), )`. If we also do detail acquisitions, the second detail in the third overview would have index `(("overview", 2), ("detail", 1))`, ...

We can get a single `MeasurementData` object, which contains lists of data, measurement parameters and hardware parameters (for each *configuration* of the measurement). The data themselves are a list of NumPy arrays for the different channels of the acquisition.

```python
# get data of a given index
measurement_data = pipeline.data[(("image", 0), )]

# get data of configuration 0, channel 0
# squeeze singleton dimensions (Inspector stacks are always 4D)
img = measurement_data.data[0][0].squeeze()
```

We can plot the image with matplotlib:

```python
from matplotlib import pyplot as plt

# NOTE: this assumes you did a 2D overview
# for 3D data, you would have to e.g., project it along the z-axis
# img = img.max(axis=0)

plt.imshow(img, cmap="magma")
```

###### Segmenting cells

To build an automation pipeline that selectively images cells in the overview with higher resolution, we first need a segmentation function that takes an image (NumPy array) and returns an integer-valued label map of the same shape.

We can use standard Python image processing functionality from libraries like `scikit-image` or `scipy`:

```python
from skimage.filters import threshold_otsu
from scipy.ndimage import gaussian_filter, label
from skimage.segmentation import clear_border
from skimage.morphology import dilation, disk


def segment(img):
    # blur and get Otsu threshold
    g = gaussian_filter(img.astype(float), 5)
    t = max(3, threshold_otsu(g))
    # label connected components, remove at border, dilate
    labels, _ = label(g > t)
    labels = clear_border(labels)
    labels = dilation(labels, disk(3))
    return labels
```

We can test our function on the image we got from the pipeline earlier & plot the results:

```python
label_map = segment(img)

plt.imshow(label_map, cmap="turbo", interpolation="nearest")
```

##### Advanced segmentation

The segmentation function does not have any specific dependencies to autoSTED, so you can use pretty much anything in the Python image processing ecosystem, e.g. deep learning-based segmentation via Cellpose [6] https://github.com/MouseLand/cellpose.

```
from cellpose.models import CellposeModel

# instantiate Cellpose model
model = CellposeModel(model_type="nuclei")

def segment_cellpose(img, diameter=30, model=model):
    # run model, return predicted instance segmentation mask
    masks, flows, styles = model.eval(
        [img],
        diameter=diameter,
        channels=[0, 0],
    )
    return masks[0]


label_map = segment_cellpose(img)
plt.imshow(label_map, cmap="turbo", interpolation="nearest")
```

##### Using the segmentation function for an overview-detail pipeline

To move from the single-level automation pipeline above to an overview-detail pipeline, we just have to add a second callback to it that will be called after each overview image and enqueue details.

Again, we can use an `AcquisitionTaskGenerator` to construct it from simple building blocks:

1. get base settings from a JSON File
2. take the location of the overview image (esp. stage position)
3. use our segmentation function wrapped in a `SegmentationWrapper` - this will apply the function to the newest image that was acquired, translate the pixel objects into scan offset & ROI size parameters.

```
from autosted.callback_buildingblocks import (
    LocationKeeper,
    NewestSettingsSelector,
)
from autosted.detection import SegmentationWrapper

# pipeline and overview generator as above
# but we now have two levels: 'overview', 'detail'
pipeline = AcquisitionPipeline(
    data_save_path="acquisition_data/test",
    hierarchy_levels=["overview", "detail"],
)
```

```python
overview_config = "config_json/test_overview.json"
next_overview_generator = AcquisitionTaskGenerator(
    "overview",
    JSONSettingsLoader(overview_config),
    SpiralOffsetGenerator(
        move_size=[50e-6, 50e-6],
        start_position=get_current_stage_coords(),
    ),
)

# acquisition task generator for details as described above
detail_config = "config_json/test_detail.json"
detail_generator = AcquisitionTaskGenerator(
    "detail",
    # 1. base settings from file
    LocationRemover(JSONSettingsLoader(detail_config)),
    # 2. locations (stage) from previous (overview) image
    LocationKeeper(NewestSettingsSelector()),
    # 3. segmentation wrapper around segmentation function
    SegmentationWrapper(segment),
)

pipeline.add_callback(next_overview_generator, "overview")
pipeline.add_callback(detail_generator, "overview")

pipeline.add_stopping_condition(
    # instead of a maximum of total images
    # we can also specify a maximum per level
    MaximumAcquisitionsStoppingCriterion(
        max_acquisitions_per_level={"overview": 5, "detail": 20}
    )
)

pipeline.run(initial_callback=next_overview_generator)
```

###### SegmentationWrapper Internals

The SegmentationWrapper object takes care of getting image data as NumPy array(s), passing it to our segmentation function and translating the pixel results back to physical microscope parameters (scan offsets, FOV lengths) and wrapping them as parameter `dicts`.

We can adjust the behavious via the constructor, e.g.:

- how to get the measurement data in which to perform detection (by default, we select the newest image of the level at which the callback is attached)
- which configuration(s) and channel(s) to use
- whether to plot results
- wheter to return scan or stage offsets
- whether to return a ready-to-use parameter dictionary (default) or just a list

of bounding boxes, which may be isnteresting if we want to e.g., manually add an offset. In the latter case, we can nest the SegmentationWrapper in a `ScanOffsetsSettingsGenerator` or `StageOffsetsSettingsGenerator` to wrap the results into a parameter dict.

```
from autosted.callback_buildingblocks import (
    ScanOffsetsSettingsGenerator,
    NewestDataSelector,
)

detail_generator = AcquisitionTaskGenerator(
    "detail",
    LocationRemover(
        JSONSettingsLoader("config_json/test2color_detail.json")
    ),
    LocationKeeper(NewestSettingsSelector()),
    ScanOffsetsSettingsGenerator(
        SegmentationWrapper(
            segment,
            data_source_callback=NewestDataSelector(
                pipeline=pipeline, level="overview"
            ),
            configurations=0,
            channels=0,
            offset_parameters="scan",
            plot_detections=True,
            return_parameter_dict=False,
        )
    ),
)
```

##### Adding on-the-fly stitching of overviews

The modular nature of autoSTED allows for easy exchange of building blocks, e.g. the data source callback of our segmentation wrapper could be exchanged for a `StitchedNewestDataSelector` to virtually stitch overview images and thus prevent skipping of cells on the border of overview tiles.

One issue that might arise here is that since the cell detector "sees the same overview multiple times" cells might be selected for detailled imagin multiple times. To prevent this, we have an `AlreadyImagedFOVFilter` that can be attached to the acquisition task generator to skip already imaged FOVs.

```
from autosted.task_filtering import AlreadyImagedFOVFilter
from autosted.callback_buildingblocks import (
    StitchedNewestDataSelector,
)

pipeline = AcquisitionPipeline(
    data_save_path="acquisition_data/test",
    hierarchy_levels=["overview", "detail"],
```

```python
)

overview_config = "config_json/test2color_overview.json"
next_overview_generator = AcquisitionTaskGenerator(
    "overview",
    JSONSettingsLoader(overview_config),
    SpiralOffsetGenerator(
        # NOTE: smaller move size to have overlap
        move_size=[40e-6, 40e-6],
        start_position=get_current_stage_coords(),
    ),
)

detail_config = "config_json/test2color_detail.json"
detail_generator = AcquisitionTaskGenerator(
    "detail",
    LocationRemover(JSONSettingsLoader(detail_config)),
    LocationKeeper(NewestSettingsSelector(pipeline, "overview")),
    SegmentationWrapper(
        segment,
        # instead of default NewestDataSelector
        # we use StitchedNewestDataSelector
        # which returns a virtually stitched image
        # of the most recent overview and its neighbors
        data_source_callback=StitchedNewestDataSelector(
            pipeline,
            "overview",
            register_tiles=False,
            offset_parameters="scan",
        ),
        offset_parameters="scan",
    ),
)
# we add a task filter
# to ignore FOVs already imaged at "detail" level
detail_generator.add_task_filters(
    AlreadyImagedFOVFilter(pipeline, "detail", 0.5, True)
)


pipeline.add_callback(next_overview_generator, "overview")
pipeline.add_callback(detail_generator, "overview")

pipeline.add_stopping_condition(
    MaximumAcquisitionsStoppingCriterion(
        max_acquisitions_per_level={"overview": 5, "detail": 20}
    )
)
```

```
pipeline.run(initial_callback=next_overview_generator)
```

##### Other detectors

Similar to how you can change the data selection callback, you can also swap the whole `SegmentationWrapper` for wrappers around functions that return a list of ROI bounding boxes or that detect just (center) coordinates.

We also provide a few "legacy" detector classes that we used over the years for detection of FISH spot (pairs) or nuclei which you can use without supplying your own detection function (see `overview_detail_nuclei_legacy.ipynb` or `overview_detail_spot_pair.ipynb`).

```
# Wrapper for custom detection functions
# that return list of bounding boxes
from autosted.detection import ROIDetectorWrapper

# Wrap custom detection function that returns
# coordinates of objects of interest (e.g. FISH spots)
from autosted.detection import CoordinateDetectorWrapper


# examples for legacy spot (pair) detectors
from autosted.detection.legacy import (
    LegacySpotPairFinder,
    SimpleSingleChannelSpotDetector,
)

# legacy nucleus (midplane) detectors
from autosted.detection.legacy import (
    SimpleNucleusMidplaneDetector,
    CellposeNucleusMidplaneDetector,
)
```
